# Platelet Bioenergetic Profiling Reveals Non-Mitochondrial Dysfunction as a Potential Biomarker of Diabetic Complications

**DOI:** 10.1101/2025.10.22.683779

**Authors:** Abdulbaki Agbas, Patricia Kluding, Mamatha Pasnoor, Whitney Shae, Jessica Sage

**Affiliations:** Kansas City University; University of Kansas Medical Center; Boehringer Ingelheim Animal Health USA Inc.

## Abstract

Diabetes is associated with systemic bioenergetic dysfunction that contributes to complications such as diabetic peripheral neuropathy (DPN). While mitochondrial impairment has been implicated, the role of non-mitochondrial pathways is less clearly defined. To address this gap, we optimized platelet-based extracellular flux assays as a minimally invasive platform to study human bioenergetics. A seeding density of 20 × 10^6^ platelets per well provided reliable respiratory measurements, consistent with the presence of 5–8 mitochondria per platelet.

Platelets from both type 2 diabetes (T2D) and DPN patients demonstrated a significant reduction in non-mitochondrial oxygen consumption rate (OCR) compared with controls. This finding suggests impairment of oxygen-consuming processes beyond the mitochondria, likely reflecting diminished activity of oxidoreductase enzymes such as NADPH oxidases, cyclooxygenases, and lipoxygenases, which regulate redox balance, inflammatory signaling, and vascular tone. In contrast, basal and ATP-linked OCR were only marginally reduced in T2D and not significantly altered in DPN, indicating that mitochondrial-linked dysfunction may be more subtle or heterogeneous in these patient populations. The consistent decrease in non-mitochondrial OCR across both T2D and DPN highlights its potential as an early and sensitive indicator of systemic bioenergetic dysregulation in diabetes.

These findings demonstrate that platelets are a practical and repeatable resource for assessing bioenergetics in humans. Diabetes impairs both mitochondrial and non-mitochondrial respiration, with the latter showing the most robust alterations. Reduced non-mitochondrial OCR may contribute independently to the pathogenesis of DPN and holds promise as a biomarker for prognosis and therapeutic monitoring in diabetic complications.

## 1. INTRODUCTION

Diabetes is the most common cause of neuropathy in the United States, with approximately 50% of diabetic patients developing some form of neuropathy over time ^1-4^. Among these, diabetic peripheral neuropathy (DPN) is one of the most prevalent and debilitating complications^4, 5^. Numerous studies have established that oxidative stress-induced tissue injury plays a central role in the pathogenesis of DPN^6-10^. A major contributing mechanism is the generation of free radicals resulting from increased glycolytic flux, which in turn leads to mitochondrial damage and impaired cellular function^11-14^.

This oxidative stress model strongly implicates mitochondrial dysfunction in the development and progression of DPN. Experimental hyperglycemia has been shown to induce mitochondrial alterations that trigger programmed cell death (apoptosis), suggesting that apoptosis-associated mitochondrial dysfunction contributes to diabetic sensory neuropathy^15-19^.

Despite this mechanistic insight, the functional status of mitochondria in human tissues, particularly in the context of diabetic neuropathy, is not well defined largely due to limited tissue accessibility. Most insulin-responsive tissues, such as skeletal muscle or adipose tissue, require invasive procedures to obtain, and even when accessible, may not reliably reflect mitochondrial dysfunction in the peripheral nervous system. This presents a significant barrier to longitudinal studies and the development of non-invasive biomarkers for mitochondrial health in diabetes.

In this context, human platelets offer a unique and practical alternative. Platelets are readily obtainable from peripheral blood with minimal invasiveness, can be sampled repeatedly, and notably, contain functional mitochondria. These characteristics make platelets an attractive and accessible surrogate tissue to evaluate mitochondrial function in diabetic patients.

The overarching premise of this study is that reduced mitochondrial function can be quantified through functional platelet mitochondria respiration. This approach has the potential to serve as a prognostic tool for T2D and DPN and to help monitor the effectiveness of therapeutic interventions. Comparing platelet mitochondrial parameters in T2D and DPN may provide a minimally invasive biomarker for prognosis of DPN and serve as a tool to monitor the response to therapeutic interventions.

In this preliminary study, we tested mitochondria respiration flux in platelets isolated blood from diabetic patients with neuropathy and compared to age-matched healthy control subjects. The assessment of metabolic function in cells isolated from human blood for treatment and diagnosis of disease is a new and important area of translational research^20^. The overall strategy of this proposal is to establish and optimize an assay method by which intact mitochondria respiration can be accessed via freshly isolated platelet.

## 2. MATERIALS AND METHODS

### 2.1. Patient recruitment

In this pilot study, we recruited both men and women between 20 and 60 years who speak English as primary language and are from any race or ethnic group. Participants will be excluded under the following conditions: **(i)** they have other medical conditions (aside from diabetes or neuropathy) that might interfere with mitochondrial function, **(ii)** they report diagnosis of pre-diabetes (to avoid confounding comparison between diabetic and control participants), **(iii)** they have peripheral neuropathy without a diagnosis of diabetes, or **(iv)** they are unable to understand or cooperate with testing procedures.

Potential research participants were recruited for this study through flyers posted in the community, information sent electronically via email and posted on websites (e.g. www.PioneersResearch.org), and trial registration on www.ClinicalTrials.gov. We also utilized the Frontiers Research Participant Registry and the Pioneers Community Research Participant Registry, following approval from the Recruitment Registry Request Committee (RRRC) at the University of Kansas Medical Center (KUMC). Potentially qualified individuals in these registry programs were identified through a search of ICD-9/ICD-10 codes in the electronic health records. They were contacted by the study team to determine interest and eligibility, then the study visit was scheduled.

Each participant completed all study procedures in a single session lasting approximately 1 hour in the Clinical and Translational Science Unit (CTSU) at KUMC. During this visit, informed consent was obtained, demographic and medical history information were collected, and a neuropathy screening questionnaire was completed. Participant diagnosis of type 2 diabetes was self-reported. The Michigan Neuropathy Screening Instrument (MNSI) includes a symptom questionnaire and a brief clinical exam administered by the study coordinator. A score >4 on the MNSI indicates presence of DPN and was used to classify participants as either DPN or Diabetes Mellitus (DM) only. Participants received compensation for their time and travel expenses related to study participation.

Whole blood was drawn onto ACD-containing vacutainer tubes by University of Kansas Clinical Translational Science Unit (CTSU) nursing staff and transported to another lab for platelet isolation and extraflux analysis (e.g. Seahorse analysis) within 1-1.5 hour of the sampling.

### 2.2. Platelet isolation

The platelet isolation was carried out according to procedures routinely used in our laboratory^21^. (Figure-1) Briefly, 8-9 ml of blood was collected in an ACD-containing vacutainer (yellow-top) tube and centrifuged at 200 x g for 20 min at room temperature (RT). The supernatant, platelet rich-plasma (PRP), was collected carefully by pipetting down to 0.5 cm above the buffy coat. The platelet activating factor inhibitor PGI^2^ was added to the PRP fraction at a concentration of 1 µg/ml, mixed gently, and subjected to centrifugation at 1,200 x g for 15 min at RT. The platelet pellet was resuspended in 1 ml of Seahorse analyzer (Sehaorse XF^e^24 Extraflux Analyzer) kit buffer, counted, and cell numbers were adjusted as 20 ×10^6^ platelets per well for extraflux analysis.

**Figure-1:**
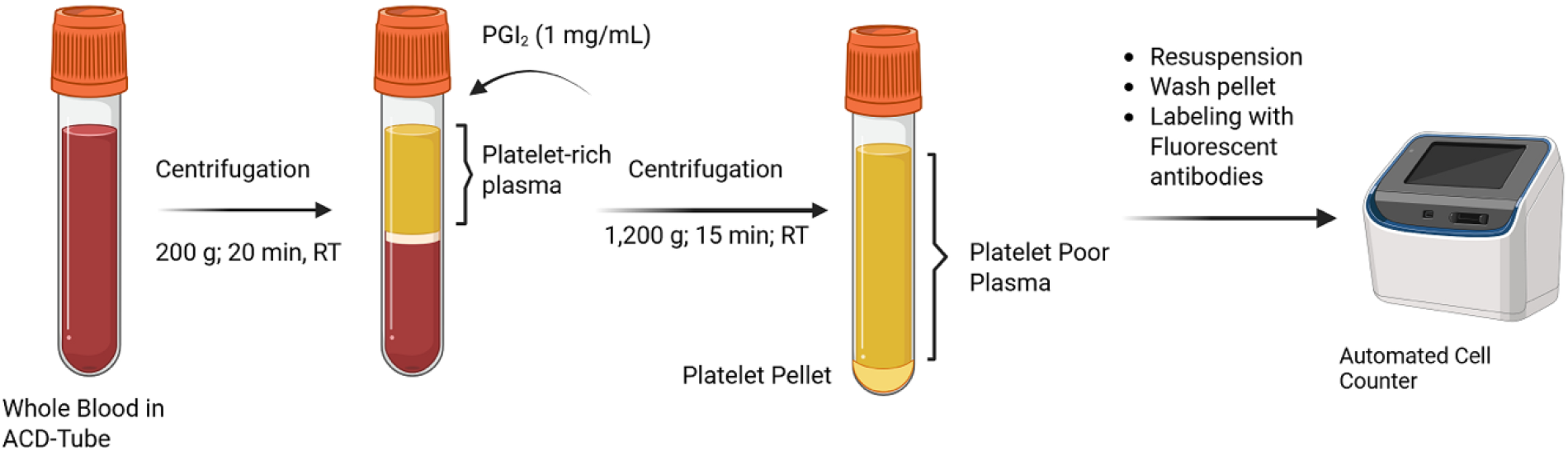
The step-wise diagram of platelet isolation from whole blood. Platelets were isolated according to our well established protocol published elsewhere^21^.

### 2.3. Cellular Bioenergetic Assays

We used XF Cell Mito Stress test kit (Agilent, Cat# 103015-100) and XF Glycolytic Rate Assay kit (Agilent Cat# 103344-100) and prepared the test plate according to manufacturer’s protocols. Assays were performed in quintuplicates to account for variation among the assay wells. Since platelets are not adhesive on the plastic substrata, we coated the cell wells with CellTac (1ug/ml), and 20×10^6^ quiescent platelets were seeded. The key cellular metabolic functions (i.e., Oxygen consumption rate (OCAR), Extracellular acidification rate (ECAR), and ATP production rate) were measured by employing the Seahorse XF^e^24 Extracellular Flux Analyzer. We collected the data and analysed by Wave software (V.2.6.4.24).

### 2.4. Statistical Analysis

To assess if whole platelet respiration can quantify reduced mitochondrial function, we tested mitochondria respiration flux in platelets isolated blood from diabetic patients with (DPN, *n* = 9) and without (T2D, *n* = 9) neuropathy against age-matched healthy controls (Control, *n* = 9). The six measures of mitochondrial respiration were Basal OCR, ATP-Linked, Proton leak, Maximal OCR, Reserved Capacity, and Non-mito OCR. We separately compared the measures of DPN and T2D patients to those of the Control group, overall conducting 12 pairwise tests using the nonparametric Mann Whitney U test. We selected a family wise error rate (FWER) of *α* = 0.05, utilizing the Holm correction for multiple testing^22^. Our hypotheses were one-sided: diabetic patients (DPN/T2D) will exhibit lower mitochondrial function as measured in platelets than age-matched healthy controls.

## 3. RESULTS

Non-mitochondrial oxygen consumption rate (OCR) was significantly reduced in both the DPN and T2D groups when compared with controls (p = 0.0053 and p = 0.0126, respectively). In contrast, basal OCR and ATP-linked respiration showed a marginally significant reduction in the T2D group relative to controls (p = 0.0588 and p = 0.0636, respectively). No significant differences in basal or ATP-linked respiration were observed between the DPN and control groups.(Fig.2) Using the Holm correction (Table-1) for multiple testing and a FWER *α* = 0.05, we conclude that Non-mito OCR is significantly lower in both the DPN group and T2D group (p-values 0.0053 and 0.0126, respectively) when compared to the Control group. We also found a marginally significant decrease in Basal OCR and ATP-Linked measures between the T2D and Control groups only (p-values 0.0588 and 0.0636, respectively (Table-2).

**Table-1:** The Holm correction for multiple testing and a family wise error rate (FWER).

| Measure | Group vs. Control | Raw p-value | Holm adjusted p-value |
| --- | --- | --- | --- |
| Basal OCR | DPN | 0.0425 | 0.3071 |
|  | T2D | 0.0059 | 0.0588 |
| ATP-Linked | DPN | 0.0384 | 0.3071 |
|  | T2D | 0.0071 | 0.0636 |
| Proton leak | DPN | 0.5184 | 0.9999 |
|  | T2D | 0.3886 | 0.9999 |
| Maximal OCR | DPN | 0.2682 | 0.9999 |
|  | T2D | 0.0425 | 0.3071 |
| Reserve Capacity | DPN | 0.3620 | 0.9999 |
|  | T2D | 0.1768 | 0.8840 |
| Non-mito OCR | DPN | 0.0004 | 0.0053 |
|  | T2D | 0.0011 | 0.0126 |

**Table-2:** Oxygen consumption rate (OCR) comparisons between diabetic groups and controls. Values represent statistical comparisons of non-mitochondrial, basal, and ATP-linked OCR between diabetic peripheral neuropathy (DPN), type 2 diabetes (T2D), and control groups. Non-mitochondrial OCR was significantly lower in both DPN and T2D groups relative to controls (p = 0.0053 and p = 0.0126, respectively). Basal OCR and ATP-linked respiration demonstrated marginal reductions in the T2D group compared with controls (p = 0.0588 and p = 0.0636, respectively). Statistical significance was defined as p < 0.05.

**Table-1:** The Holm correction for multiple testing and a family wise error rate (FWER).
| OCR Measure | Group Comparison | p-value | Interpretation |
| --- | --- | --- | --- |
| Non-mito OCR | DPN vs. Control | 0.0053 | Significant ↓ |
|  | T2D vs. Control | 0.0126 | Significant ↓ |
| Basal OCR | T2D vs. Control | 0.0588 | Marginal ↓ (trend) |
| ATP-linked OCR | T2D vs. Control | 0.0636 | Marginal ↓ (trend) |

**FIGURE-2.**
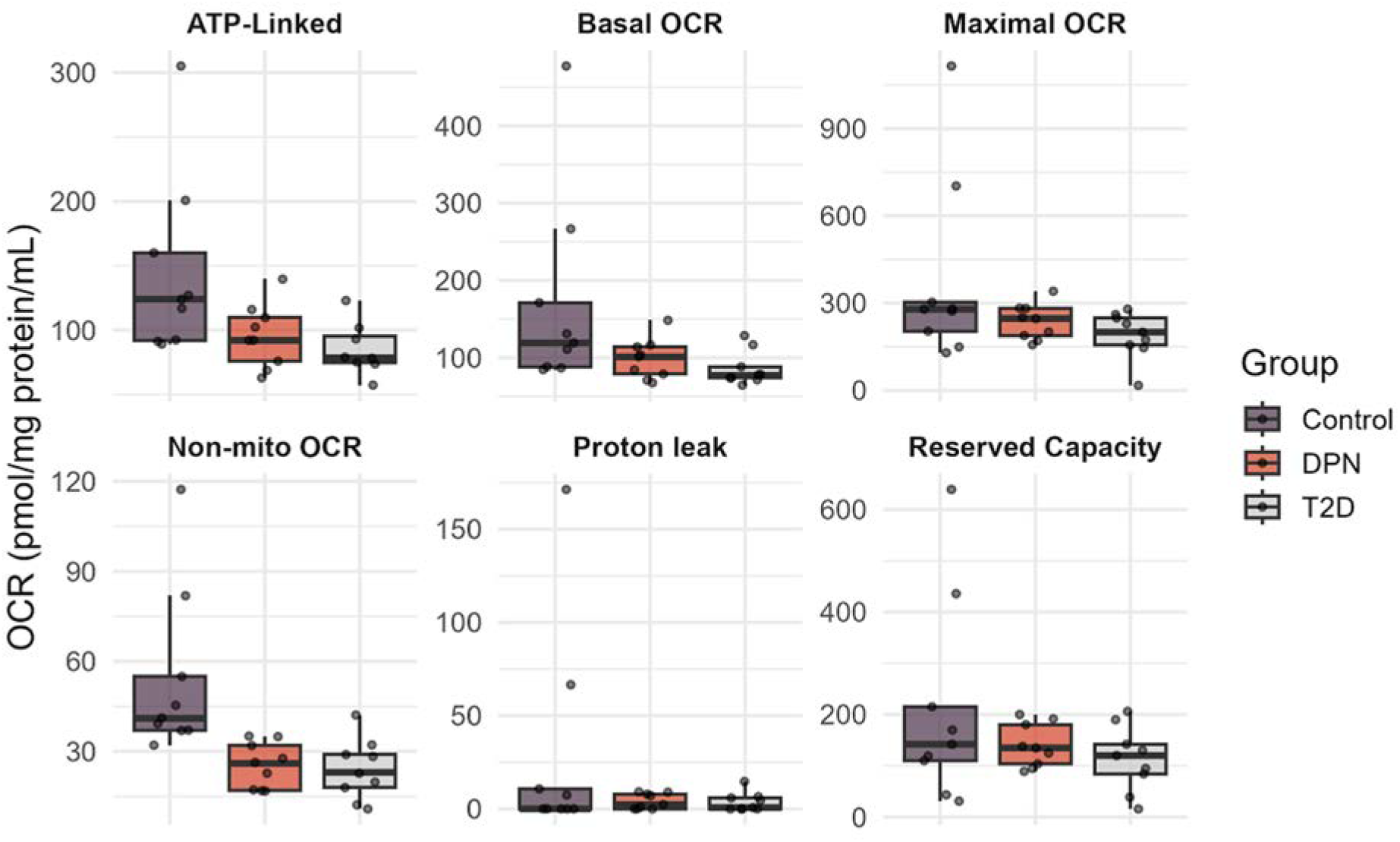
Platelet respiration in Control, T2D, and DPN groups. DPN patients showed a significant reduction only in non-mitochondrial OCR, while T2D patients exhibited lower non-mito, basal, and ATP-linked OCR compared with Controls. No other respiratory parameters differed significantly between groups.

Overall, when comparing patients with DPN to Control patients, only Non-mito OCR was significantly lower for DPN patients. Type 2 Diabetic patients without peripheral neuropathy (T2D) had Non-mito OCR, Basal OCR, and ATP-Linked measures at least marginally lower than Control patients. We have insufficient evidence to conclude that any other measures are significantly different between diabetic patients and age-matched controls, as quantified through whole platelet respiration.

## 4. DISCUSSION

The platelet mitochondrial extraflux analyses showed that platelets are practical resource for studying bioenergetics in human with repetitive sampling with minimal invasion. 20×10^6^ platelets per extraflux cell plate well is optimized in order to provide sufficient mitochondria as each platelets have about 5-8 mitochondria^23, 24^.

The significant reduction in non-mitochondrial OCR observed in both T2D and DPN groups suggests that diabetes broadly affects oxygen-consuming processes beyond the mitochondria (sup.Fig 1-A), such as oxidase-mediated pathways, while not a noticeable changes observed in extracellular acidification rate (ECAR) (sup. Fig.1B and 1C). This finding indicates a systemic shift in cellular bioenergetics that is not limited to mitochondrial respiration and may reflect altered redox balance or diminished activity of extra-mitochondrial enzymes in diabetic patients.

The marginal reductions in basal OCR and ATP-linked respiration detected in the T2D group further suggest that mitochondrial function itself is compromised, although these findings did not reach conventional levels of statistical significance. This trend is consistent with reviews and prior reports of diminished mitochondrial efficiency and ATP production in type 2 diabetes^25-28^. The absence of significant differences between DPN and control groups for these parameters may be due to sample size limitations or heterogeneity in the neuropathy population, and warrants further investigation with larger cohorts.

The observed impairment of non-mitochondrial oxygen consumption in both T2D and DPN groups underscores the importance of extra-mitochondrial bioenergetic pathways in the pathophysiology of diabetes^29^. Non-mitochondrial OCR reflects the activity of oxidoreductase enzymes such as NADPH oxidases^30-33^, cyclooxygenases^34^, and lipoxygenases^35^, which play key roles in maintaining redox balance, regulating inflammatory responses, and modulating vascular tone (Fig.3).

**Figure-3.**
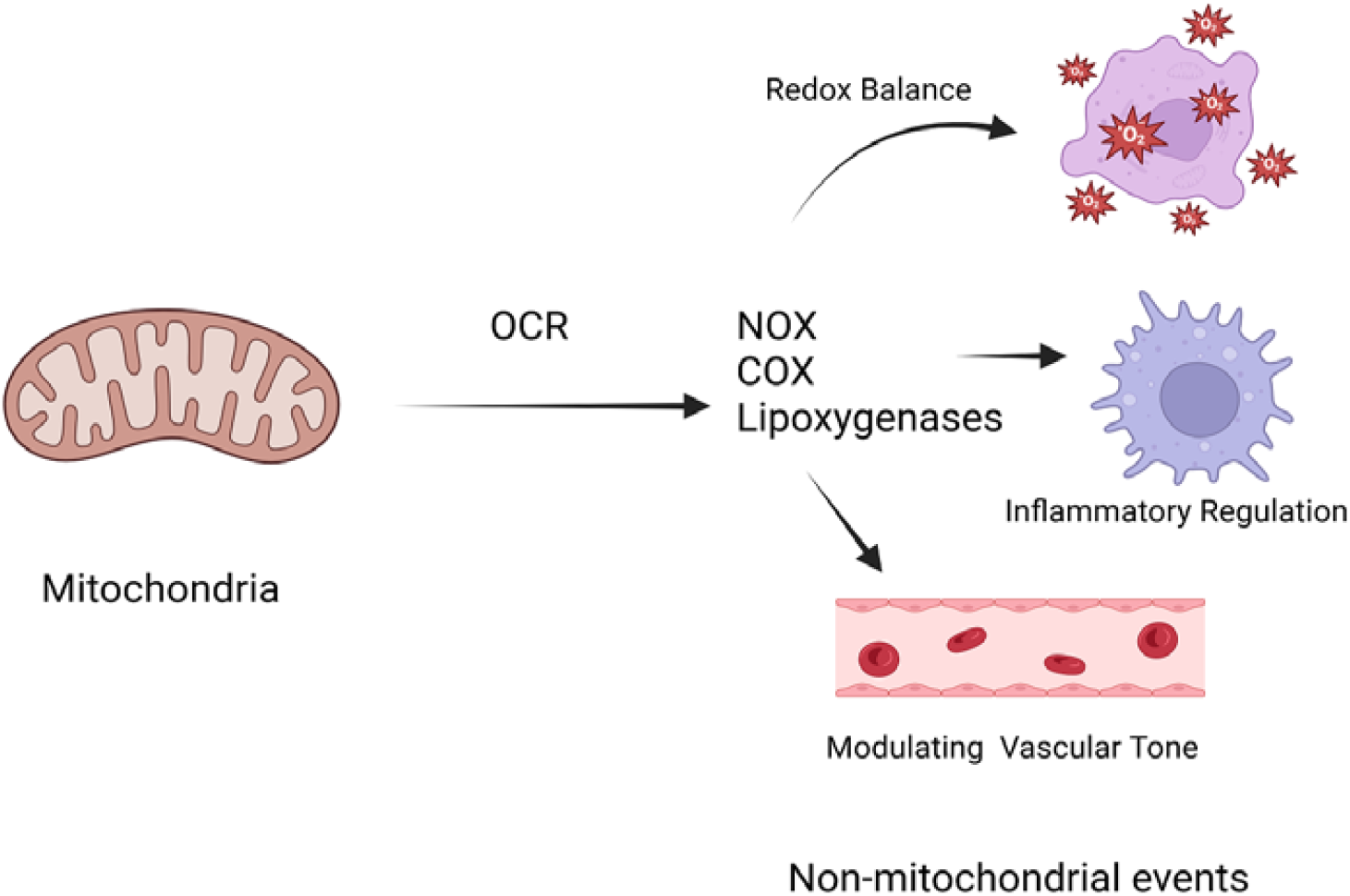
Non-mitochondrial oxygen consumption rate (OCR) affects several biological events.

A reduction in this parameter suggests diminished capacity for redox signaling and altered inflammatory activity, processes that are central to the development of diabetic complications including neuropathy. The consistent decrease in non-mitochondrial oxygen consumption rate (OCR)^36^ observed in diabetes may reflect early, extra-mitochondrial bioenergetic dysregulation^29^, driven by upregulated oxidase pathways (e.g., NADPH oxidases, COX/LOX), and thus could serve as a sensitive blood-based marker of systemic metabolic dysfunction that is not simply downstream of mitochondrial impairment. These findings highlight that alterations in non-mitochondrial pathways may not simply be secondary to mitochondrial dysfunction but could contribute independently to the pathogenesis of DPN.

## 5. CONCLUSION

These results support the hypothesis that diabetes impairs both mitochondrial and non-mitochondrial bioenergetics, with the strongest and most consistent effects observed in non-mitochondrial respiration. Importantly, the observed alterations in platelet respiration may provide an accessible biomarker of systemic metabolic dysfunction in diabetes and its complications.

## 6. APPENDICES

### SUPPLEMENTARY FIGURES

**Sup.Fig-1A.**
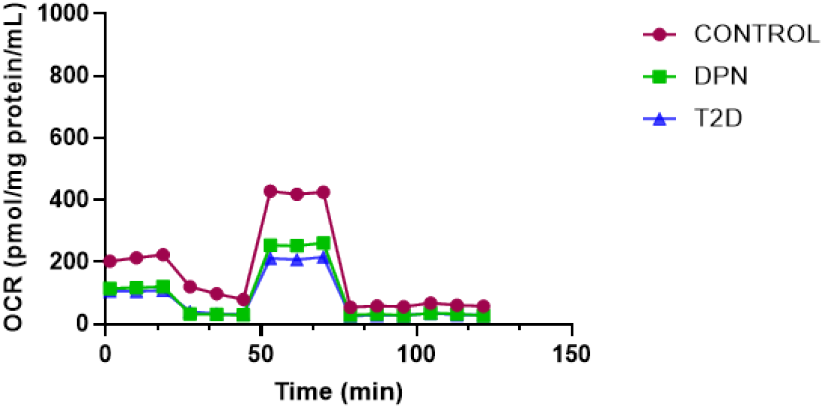
Platelet OCR profile

**Sup.Fig-1B.**
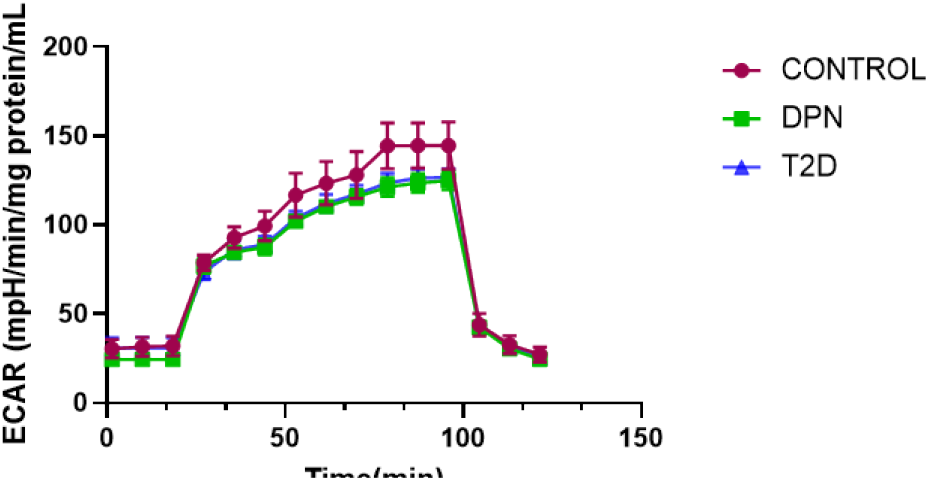
Platelet ECAR profile

**Sup.Fig-1C.**
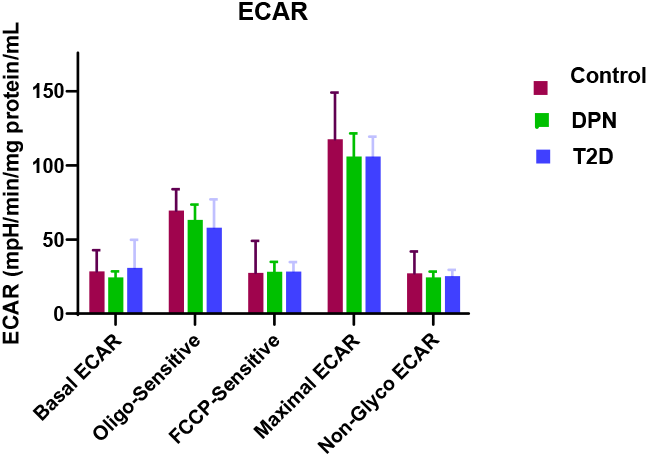
ECAR parameters by group

## REFERENCES

1. Feldman, E.L., et al., Diabetic neuropathy. Nat Rev Dis Primers, 2019. 5(1): p. 42.

2. Hicks, C.W. and E. Selvin, Epidemiology of Peripheral Neuropathy and Lower Extremity Disease in Diabetes. Curr Diab Rep, 2019. 19(10): p. 86.

3. Pop-Busui, R., et al., in Diagnosis and Treatment of Painful Diabetic Peripheral Neuropathy. 2022: Arlington (VA).

4. Sun, J., et al., Prevalence of peripheral neuropathy in patients with diabetes: A systematic review and meta-analysis. Prim Care Diabetes, 2020. 14(5): p. 435–444.

5. Smith, A.G. and J.R. Singleton, Diabetic neuropathy. Continuum (Minneap Minn), 2012. 18(1): p. 60–84.

6. Lin, Q., et al., Oxidative Stress in Diabetic Peripheral Neuropathy: Pathway and Mechanism-Based Treatment. Mol Neurobiol, 2023. 60(8): p. 4574–4594.

7. Kasznicki, J., et al., Evaluation of oxidative stress markers in pathogenesis of diabetic neuropathy. Mol Biol Rep, 2012. 39(9): p. 8669–78.

8. Obrosova, I.G., et al., Role of nitrosative stress in early neuropathy and vascular dysfunction in streptozotocin-diabetic rats. Am J Physiol Endocrinol Metab, 2007. 293(6): p. E1645–55.

9. Vincent, A.M., M. Brownlee, and J.W. Russell, Oxidative stress and programmed cell death in diabetic neuropathy. Ann N Y Acad Sci, 2002. 959: p. 368–83.

10. Vincent, A.M., et al., Oxidative stress in the pathogenesis of diabetic neuropathy. Endocr Rev, 2004. 25(4): p. 612–28.

11. Roman-Pintos, L.M., et al., Diabetic Polyneuropathy in Type 2 Diabetes Mellitus: Inflammation, Oxidative Stress, and Mitochondrial Function. J Diabetes Res, 2016. 2016: p. 3425617.

12. Stefano, G.B., S. Challenger, and R.M. Kream, Hyperglycemia-associated alterations in cellular signaling and dysregulated mitochondrial bioenergetics in human metabolic disorders. Eur J Nutr, 2016. 55(8): p. 2339–2345.

13. Hosseini, A. and M. Abdollahi, Diabetic neuropathy and oxidative stress: therapeutic perspectives. Oxid Med Cell Longev, 2013. 2013: p. 168039.

14. Figueroa-Romero, C., M. Sadidi, and E.L. Feldman, Mechanisms of disease: the oxidative stress theory of diabetic neuropathy. Rev Endocr Metab Disord, 2008. 9(4): p. 301–14.

15. Otero, M.G., et al., Hyperglycemia-induced mitochondrial abnormalities in autonomic neurons via the RAGE axis. Sci Rep, 2025. 15(1): p. 25231.

16. Zhang, Z., et al., The impact of oxidative stress-induced mitochondrial dysfunction on diabetic microvascular complications. Front Endocrinol (Lausanne), 2023. 14: p. 1112363.

17. Zhang, L., et al., Hyperglycemia alters the schwann cell mitochondrial proteome and decreases coupled respiration in the absence of superoxide production. J Proteome Res, 2010. 9(1): p. 458–71.

18. Vincent, A.M., et al., Short-term hyperglycemia produces oxidative damage and apoptosis in neurons. FASEB J, 2005. 19(6): p. 638–40.

19. Srinivasan, S., M. Stevens, and J.W. Wiley, Diabetic peripheral neuropathy: evidence for apoptosis and associated mitochondrial dysfunction. Diabetes, 2000. 49(11): p. 1932–8.

20. Kramer, P.A., et al., A review of the mitochondrial and glycolytic metabolism in human platelets and leukocytes: implications for their use as bioenergetic biomarkers. Redox Biol, 2014. 2: p. 206–10.

21. Luthi-Carter, R., et al., Location and function of TDP-43 in platelets, alterations in neurodegenerative diseases and arising considerations for current plasma biobank protocols. Sci Rep, 2024. 14(1): p. 21837.

22. Aickin, M. and H. Gensler, Adjusting for multiple testing when reporting research results: the Bonferroni vs Holm methods. Am J Public Health, 1996. 86(5): p. 726–8.

23. Shehwar, D., et al., Platelets and mitochondria: the calcium connection. Mol Biol Rep, 2025. 52(1): p. 276.

24. Melchinger, H., et al., Role of Platelet Mitochondria: Life in a Nucleus-Free Zone. Front Cardiovasc Med, 2019. 6: p. 153.

25. Crescenzo, R., et al., Mitochondrial efficiency and insulin resistance. Front Physiol, 2014. 5: p. 512.

26. Donnell, R.A., J.E. Carre, and C. Affourtit, Acute bioenergetic insulin sensitivity of skeletal muscle cells: ATP-demand-provoked glycolysis contributes to stimulation of ATP supply. Biochem Biophys Rep, 2022. 30: p. 101274.

27. Ruegsegger, G.N., et al., Altered mitochondrial function in insulin-deficient and insulin-resistant states. J Clin Invest, 2018. 128(9): p. 3671–3681.

28. Weksler-Zangen, S., Is Type 2 Diabetes a Primary Mitochondrial Disorder? Cells, 2022. 11(10).

29. Avila, C., et al., Platelet mitochondrial dysfunction is evident in type 2 diabetes in association with modifications of mitochondrial anti-oxidant stress proteins. Exp Clin Endocrinol Diabetes, 2012. 120(4): p. 248–51.

30. Kallenborn-Gerhardt, W., K. Schroder, and A. Schmidtko, NADPH Oxidases in Pain Processing. Antioxidants (Basel), 2022. 11(6).

31. Eid, S.A., et al., Targeting the NADPH Oxidase-4 and Liver X Receptor Pathway Preserves Schwann Cell Integrity in Diabetic Mice. Diabetes, 2020. 69(3): p. 448–464.

32. Eid, S.A., et al., Nox, Nox, Are You There? The Role of NADPH Oxidases in the Peripheral Nervous System. Antioxid Redox Signal, 2022. 37(7-9): p. 613–630.

33. Olukman, M., et al., Treatment with NADPH oxidase inhibitor apocynin alleviates diabetic neuropathic pain in rats. Neural Regen Res, 2018. 13(9): p. 1657–1664.

34. Kellogg, A.P., et al., Protective effects of cyclooxygenase-2 gene inactivation against peripheral nerve dysfunction and intraepidermal nerve fiber loss in experimental diabetes. Diabetes, 2007. 56(12): p. 2997–3005.

35. Stavniichuk, R., et al., Role of 12/15-lipoxygenase in nitrosative stress and peripheral prediabetic and diabetic neuropathies. Free Radic Biol Med, 2010. 49(6): p. 1036–45.

36. Chacko, B.K., et al., The Bioenergetic Health Index: a new concept in mitochondrial translational research. Clin Sci (Lond), 2014. 127(6): p. 367–73.

